# Automated high throughput animal DNA metabarcode classification

**DOI:** 10.1101/219675

**Authors:** Teresita M. Porter, Mehrdad Hajibabaei

## Abstract

Until now, there has been difficulty assigning names to animal barcode sequences isolated directly from eDNA in a rapid, high-throughput manner, providing a measure of confidence for each assignment. To address this gap, we have compiled nearly 1 million marker gene DNA barcode sequences appropriate for classifying chordates, arthropods, and flag members of other major eukaryote groups. We show that the RDP naïve Bayesian classifier can assign the same number of queries 19 times faster than the popular BLAST top hit method and reduce the false positive rate by two-thirds. As reference databases become more representative of current species diversity, confidence in taxonomic assignments should continue to improve. We recommend that investigators can improve the performance of species-level assignments immediately by supplementing existing reference databases with full-length DNA barcode sequences from representatives of local fauna.

## Introduction

Any ecological investigation such as environmental biomonitoring requires the identification of individual specimens by comparing morphological characters in specimens with those in taxonomic keys. The ‘taxonomic impediment’ describes how traditional methods may be constrained by the time it takes to process large numbers of individuals, lack of taxonomic expertise and taxonomic keys, as well as difficulties in identifying partial or immature specimens that lack the appropriate morphological characters for identification^1^. The DNA metabarcoding approach is highly scalable, capable of surveying bulk environmental samples (e.g. soil, water, passively collected biomass) in a high throughput manner, essentially shifting the taxonomic assignment of organisms from individual taxonomic experts to computational algorithms that can put a name on an anonymous DNA sequence based on comparisons to a reference sequence library ^2^. For truly high throughput biomonitoring, concurrent advances in automated assignment methods are needed to keep pace with advances in DNA sequencing throughput.

The Ribosomal Database Project (RDP) classifier uses a naïve Bayesian approach to make taxonomic assignments based on a reference dataset ^3^. The original classifier was developed using a prokaryote 16S ribosomal DNA (rDNA) reference set. The classifier can be trained, however, to classify taxa using any DNA marker. For example, using the ITS or LSU rDNA regions the method has been used to taxonomically assign fungal sequences ^4^. One advantage of using the RDP classifier over the more widely used top BLAST hit method is speed. This method is much faster and can process large datasets from high throughput sequencing in a fraction of the time that it would take with BLAST. Additionally, unlike BLAST, the RDP classifier was specifically developed to make assignments to a variety of ranks and provides a measure of confidence for each assignment at each taxonomic rank.

Extensive reference databases and tools already exist for prokaryotes and fungi (UNITE, RDP) ^4–10^. However, most of the DNA metabarcode papers for animal species have used various iterations of BLAST for classification of CO1 sequences from bulk environmental samples (e.g. Gibson et al. 2015). Although the BOLD database contains a reference set of CO1 barcode sequences ^11^, it does not provide support for analysis of large batches of CO1 metabarcodes generated by high throughput sequencing as this system was designed more as a curation and analysis tool for individual specimens. There is a training set to classify Insecta CO1 sequences using the RDP classifier but it cannot be used to identify the broader range of eukaryotes targeted with the CO1 marker from complex environmental DNA (eDNA) samples ^12^.

The purpose of this study is to take advantage of naïve Bayesian approach for high throughput assignment of sequences generated from animal DNA metabarcoding studies. By providing confidence scores at each rank, using a method that is both open-source and well-documented, we believe this approach can set the stage for application of DNA metabarcoding in large-scale real-world biomonitoring scenarios. (1) We compiled a comprehensive training set for the RDP classifier focusing on the Chordata and Arthropoda, the two largest groups of publically available CO1 sequences. (2) We benchmark the performance of the classifier for different CO1 sequence lengths, taxonomic groups, with a focus on taxa of particular importance for freshwater biomonitoring. (3) We provide guidelines for bootstrap support cutoffs to reduce false positive taxonomic assignments. (4) We show that the RDP classifier makes more taxonomic assignments per minute than the top BLAST hit approach and has lower false positive rates (FPR).

## Results

The taxonomic composition of the CO1 Eukaryote v1 training set is summarized in Table 1 and in detail in Table S1. A similar summary table is shown for the CO1 Eukaryote v2 training set in Table S2. Outgroup taxa were also included to help sort non-Arthropod and non-Chordata taxa present in eDNA samples into broad groups such as fungi, diatoms, or nematodes.

**Table 1:**
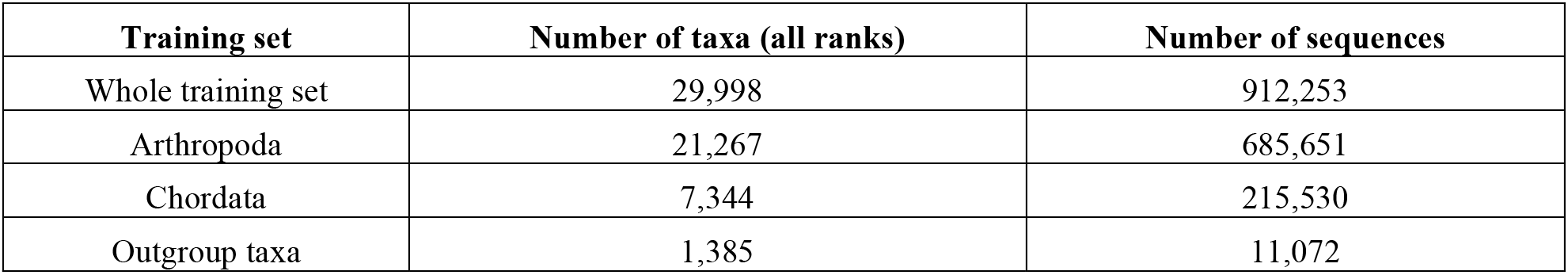
CO1 Eukaryote v1 set summary.

The proportion of singletons in the dataset can indicate groups with low taxonomic sampling coverage in the reference set and so is summarized for both the v1 and v2 training sets in Table S3. The proportion of singleton genera in the genus-trained classifier is 23% compared with the proportion of singleton species in the species-trained classifier at 33%. Since the classifier is not meant to classify taxa not represented in the training set, this means that nearly a quarter of the sequences in the CO1 Eukaryote v1 training set are not included in the leave-one-out testing which is in turn used to assess bootstrap support cutoff levels. In this study we define a false positive (FP) as an incorrect taxonomic assignment with a bootstrap support value greater than the cutoff. To avoid making FP assignments, the bootstrap cutoffs presented in this study should be treated as *minimum* cutoff values. Taxa for the training set were sampled to emphasize Arthropoda and Chordata since these were the best-represented eukaryote phyla in the GenBank nucleotide database. Figure 1 shows the proportion of correctly assigned sequences for a variety of query lengths at a variety of taxonomic ranks. Since the classifier is not meant to classify taxa not represented in the database, leave-one-out testing results from singletons were excluded from the results and no bootstrap support cutoff was used. Classifier accuracy is highest at more inclusive taxonomic ranks, especially for fragments 200bp or longer.

**Figure 1:**
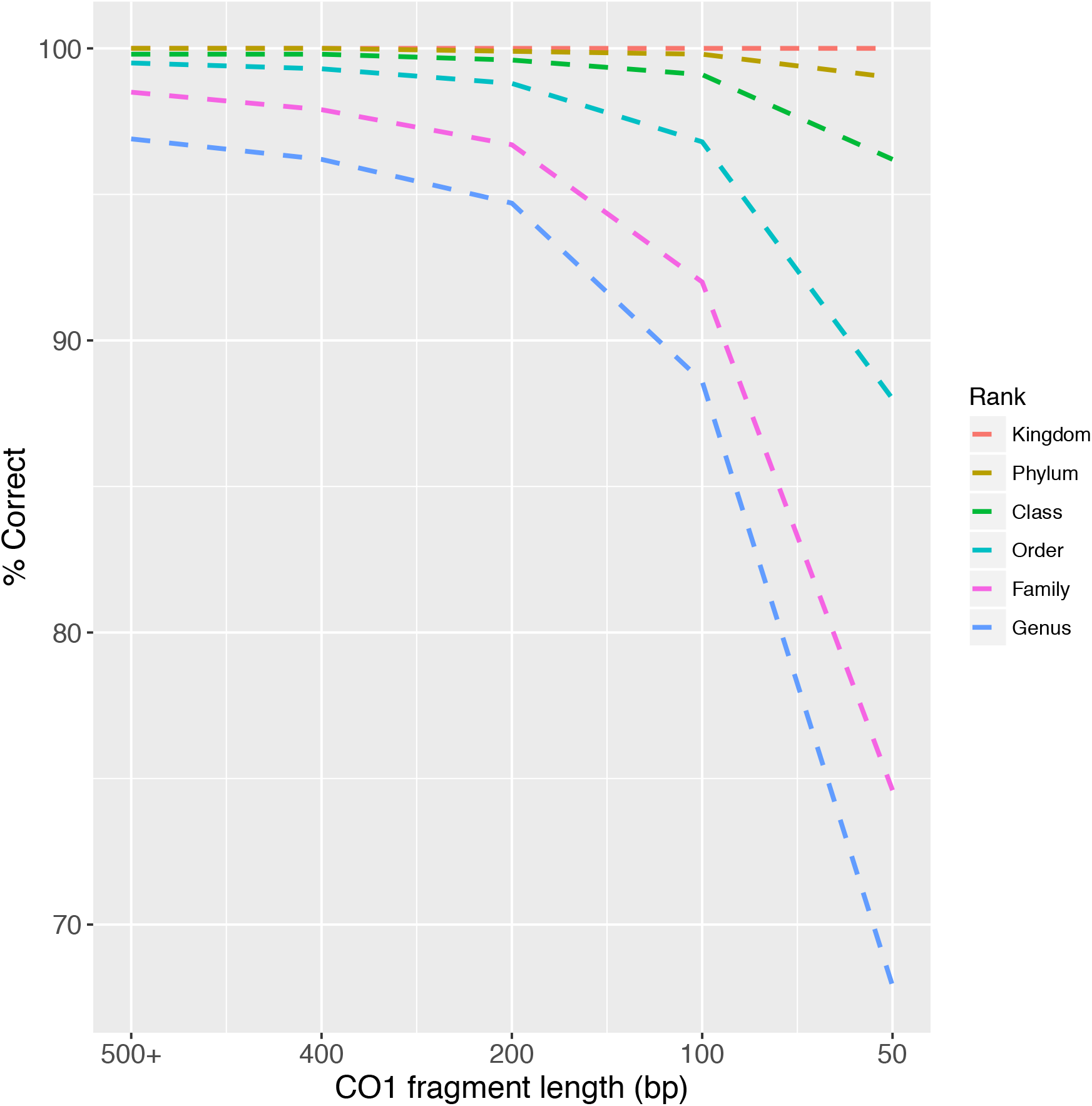
Proportion of correct taxonomic assignments increases with more inclusive taxonomic ranks and longer CO1 sequences. Results summarize the proportion of correctly assigned sequences during leave-one-out testing of the CO1 Eukaryote v1 training set.

A receiver operator characteristic (ROC) curve shows the relationship between the false positive rate (FPR) and true positive rate (TPR). This is calculated from the leave-one-out testing results of full length (500 bp+) sequences as the bootstrap support cutoff is tuned from 0 to 100. Results from singletons were not included. The FPR represents the proportion of incorrect assignments with a high bootstrap support value out of all incorrect assignments. The TPR represents the proportion of correct assignments with a high bootstrap support value out of all correct assignments. The ROC curves for full length CO1 barcode sequences to various taxonomic ranks, and for a range of fragment lengths at the genus rank is shown in Figure S1. Points lying above the 50% line indicate results better than those obtained by chance. The high area-under-the-curve values indicate high true positive rates and good classifier performance across a wide range of bootstrap support values. For full length CO1 sequences, the TPR at all taxonomic ranks is high indicating that most of the assignments are correctly assigned. For the shortest CO1 sequences assigned to the genus rank, the TPR increases with higher bootstrap support cutoffs. The FPR also increases as the bootstrap support cutoff increases from left to right, indicating that the relative number of incorrectly assigned sequences from FPs increases as true negatives (TNs) are filtered out.

Since classification performance varies with fragment size and taxonomic assignment rank, we have calculated a matrix of minimum bootstrap support value cutoffs to obtain 99% correct assignments assuming the query sequence is present in the training set (Table 2). Singletons were excluded from this analysis and cutoffs are based on 77% of the sequences in the original training set. Also shown is the corresponding reduction in the proportion of classified sequences after applying the minimum bootstrap support cutoff values. A similar table for the Eukaryote v2 classifier trained to the species rank is shown in Table S4. Generally as the amount of sequence information decreases with decreasing CO1 sequence length, higher bootstrap support cutoff values are needed to observe 99% correct assignments. Similarly, as assignments are made to increasingly specific ranks, higher cutoff values are required to observe 99% correct assignments.

**Table 2:** Bootstrap support cutoff values that produced at least 99% correct assignments during CO1 Eukaryote v1 leave-one-out testing.

| Rank | 500bp+ | 400bp | 200bp | 100bp | 50bp |
| --- | --- | --- | --- | --- | --- |
|  | **Minimum bootstrap support cutoff (%)** |  |  |  |  |
| Superkingdom | 0 | 0 | 0 | 0 | 0 |
| Kingdom | 0 | 0 | 0 | 0 | 0 |
| Phylum | 0 | 0 | 0 | 0 | 0 |
| Class | 0 | 0 | 0 | 0 | 60 |
| Order | 0 | 0 | 10 | 40 | 80 |
| Family | 20 | 20 | 30 | 40 | 80 |
| Genus | 70 | 60 | 60 | 60 | N/A |
|  | **Reduction of sequences classified after applying minimum bootstrap support cutoff (%)** |  |  |  |  |
| Superkingdom | 0.0 | 0.0 | 0.0 | 0.0 | 0.0 |
| Kingdom | 0.0 | 0.0 | 0.0 | 0.0 | 0.0 |
| Phylum | 0.0 | 0.0 | 0.0 | 0.0 | 0.0 |
| Class | 0.0 | 0.0 | 0.0 | 0.0 | 13.2 |
| Order | 0.0 | 0.0 | 0.2 | 5.7 | 60.7 |
| Family | 0.7 | 1.2 | 4.1 | 17.1 | 78.7 |
| Genus | 3.4 | 4.7 | 10.8 | 31.0 | N/A |

'N/A', not applicable, refers to the inability to observe 99% correct taxonomic assignments.

Applying a bootstrap support cutoff can reduce the proportion of incorrect taxonomic assignments. Including the results from singletons during leave-one-out testing provided a convenient way to simulate the taxonomic assignment of sequences without congenerics in the database. Figure S2 shows the proportion of incorrect assignments for Arthropoda sequences both with and without using bootstrap support cutoffs. The 70% bootstrap support cutoff value was selected for full length (500 bp+) CO1 sequences as shown in Table 2. When classifying Arthropoda sequences, 23% of which were known to have no congenerics in the training set (Table S3), applying a 70% bootstrap support cutoff at the genus rank reduced the misclassification rate for nearly all classes to ∽1% for most classes while reducing the number of assigned sequences by ∽ 3%. A similar analysis with Chordata is shown in Figure S3. The proportion of incorrect assignments for all phyla, Arthropoda and Chordata classes, as well as for the orders in the large Insecta and Actinopteri groups are shown in the Supplementary Material Tables S5-S9. When we focus on groups that are particularly important in freshwater biomonitoring, we see that database sequences are highly skewed towards Diptera and that although the proportion of incorrect classification varies across groups the application of a bootstrap support cutoff reduces these rates to ∽ 1% incorrect assignments (Table 3). One exception is for sequences in Megaloptera that have a higher proportion of incorrect assignments (1.7%) even after using a 70% bootstrap support cutoff at the genus rank for full length (500 bp+) CO1 sequences. These tables clearly show how database representation and misclassification rates can vary across taxonomic groups.

**Table 3:** Representation of freshwater biomonitoring taxa in the Eukaryote CO1v1 training set.

| **Class** | **Order** | **No. reference sequences** | **% Incorrect (No cutoff)** | **% Incorrect (Cutoff)** |
| --- | --- | --- | --- | --- |
| Bivalvia | - | 667 | 3.7 | 0.3 |
| Clitellata | - | N/A | N/A | N/A |
| Gastropoda | - | 1,896 | 3.7 | 0.4 |
| Insecta | Coleoptera | 89,484 | 7.5 | 1.1 |
| Insecta | Diptera | 118,896 | 3.8 | 0.8 |
| Insecta | Ephemeroptera | 6,722 | 2.8 | 0.3 |
| Insecta | Megaloptera | 469 | 3.6 | 1.7 |
| Insecta | Odonata | 3,553 | 6.9 | 1.2 |
| Insecta | Plecoptera | 2,679 | 2.7 | 0.1 |
| Insecta | Trichoptera | 17,277 | 3.1 | 0.3 |
| Malacostraca | Amphipoda | 8,483 | 3.4 | 1.3 |
| Malacostraca | Isopoda | 3,659 | 2.9 | 0.1 |
| Polychaeta | - | 888 | 2.8 | 0.2 |
| Turbellaria | - | N/A | N/A | N/A |

N/A, not applicable, as of October 2016 there are no full length CO1 sequences identified to the species rank in the GenBank nucleotide database. Where indicated, we used a genus bootstrap value of 70% as a cutoff

Classification performance may also vary for partial CO1 sequences whether they are sampled randomly from across the barcoding region (Figure 1) or if they are anchored by specific CO1 primers (Figure 2). The coverage of primer-anchored 200 bp sequences sampled from the dataset varies across the length of the barcoding region. Since primers are often trimmed before submission to GenBank, it was not surprising that the Folmer barcoding primers, and other primers designed near the 5’ and 3’ end of the barcoding region, had especially low coverage in our training set (Figure 3). The proportion of correct assignments of primer-anchored 200 bp sequences with and without 60% bootstrap support (Table 2) is also shown. Singletons were not included in this analysis. The proportion of correct assignments is especially high at the order to kingdom ranks with some variation among primers. After applying the bootstrap support cutoff, the proportion of correct taxonomic assignments rose to ∽99% across all primers at the genus, family, and order ranks.

**Figure 2:**
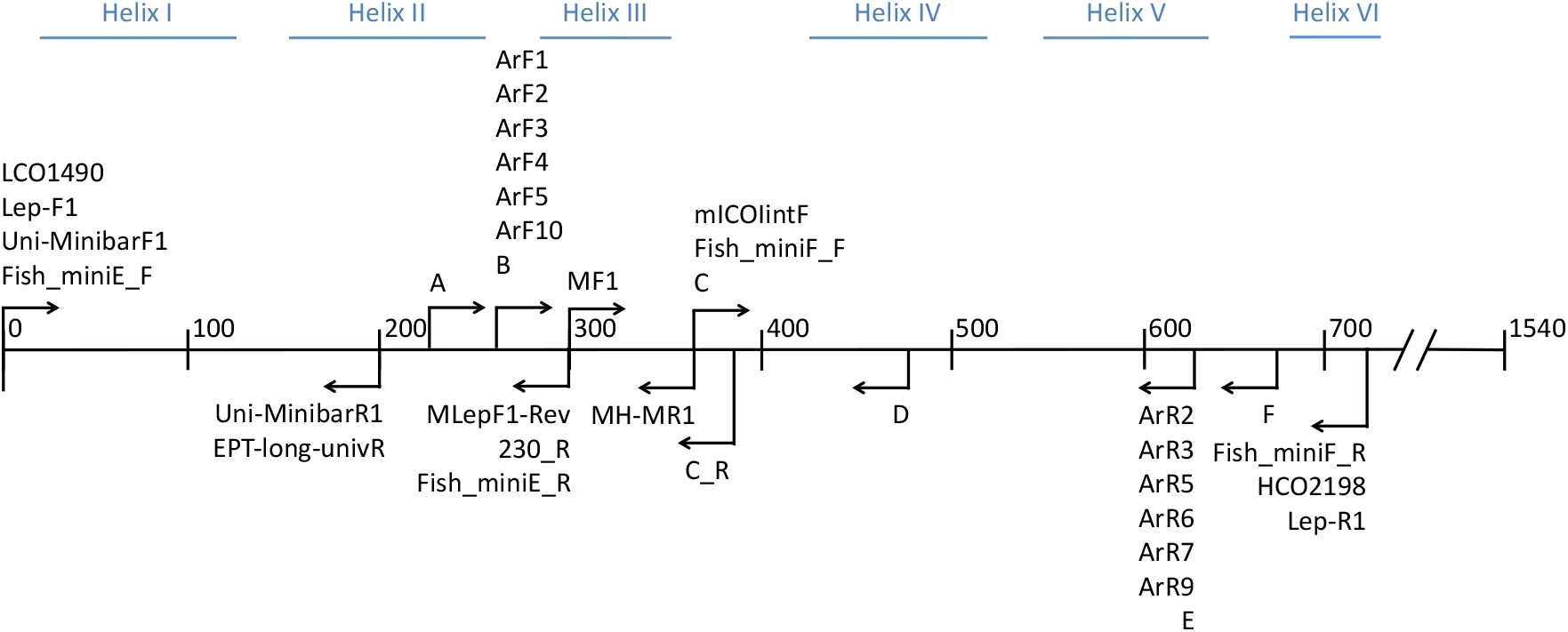
CO1 primers included in this study. Primer map of the CO1 barcoding region showing the relative position and direction of the primer-anchored 200 bp fragments analyzed in this study. The CO1 helix regions that are embedded in the mitochondrial inner membrane are also shown for reference.

**Figure 3:**
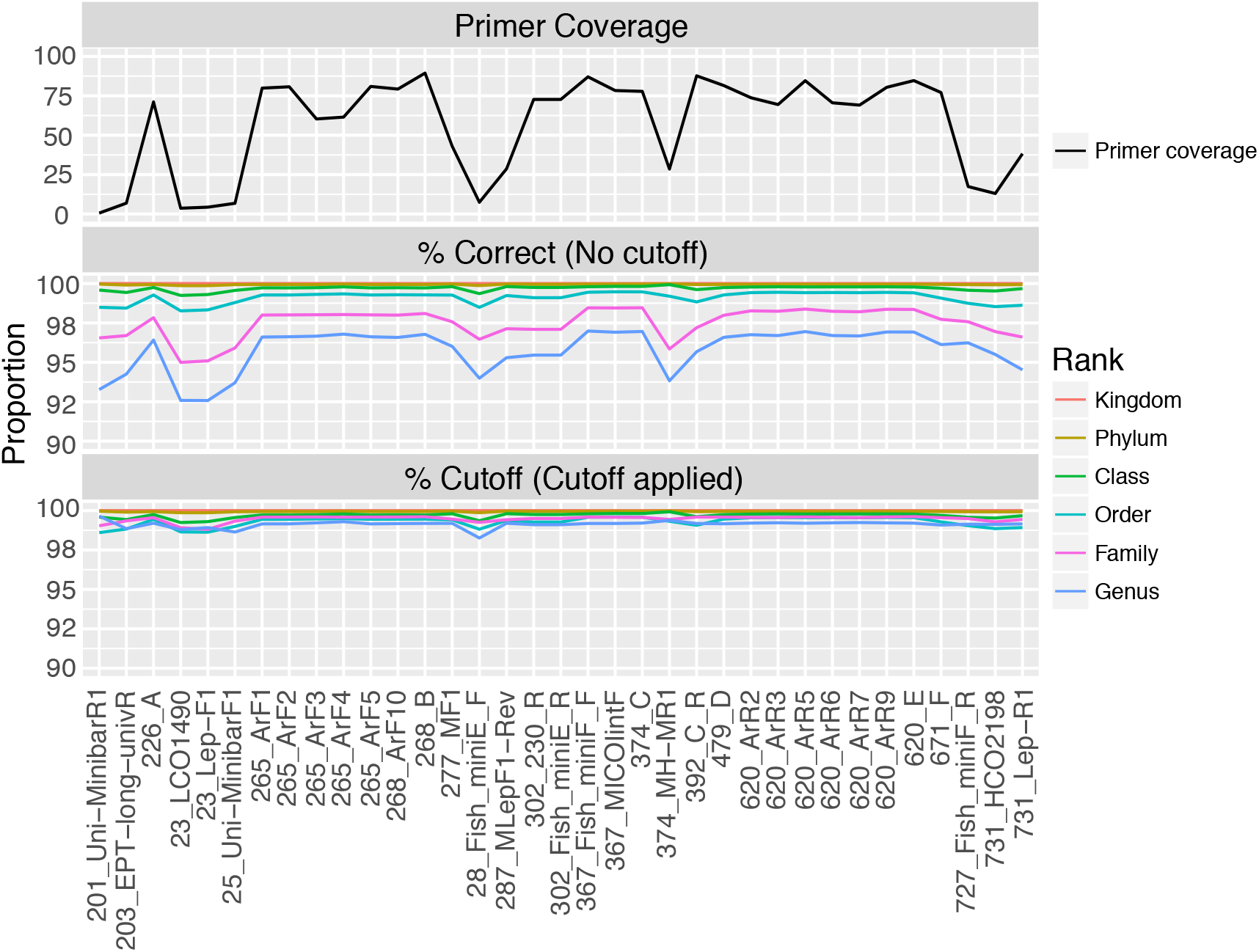
Proportion of correctly assigned primer-anchored 200 bp sequences can vary widely across the CO1 barcoding region before applying a bootstrap support cutoff. Primer names are prefixed with the outermost alignment position along the CO1 barcoding region and are arranged along the x-axis in the order that they would be encountered from the 5’ to 3’ end. Top panel: Coverage of primer-anchored 200 bp sequences in the CO1 Eukaryote v1 training set. Middle panel: Proportion of correct taxonomic assignments. Bottom panel: Proportion of correct assignments after filtering by a 60% bootstrap support cutoff at the genus rank. Note the differing limits on the y-axes.

A comparison of taxonomic assignment outcomes using the top BLAST hit method and the RDP Classifier with the CO1 Eukaryote v1 training set is shown for all primer-anchored 200 bp fragments in Table 4 and Figure S4. Using BLAST, no hits were returned for some queries because the expect value (e-value) was greater than the default cutoff of 10. In contrast, using the RDP classifier, a result was returned for every query. Assignment accuracy (Table 4) is highest for the top BLAST hit method, however, the FPR is ∽ 3 times higher for BLAST than for the RDP classifier. This is significant because in this example, 397,820 taxonomic assignments are classified as ‘good’ based on the top BLAST hit metrics but they are actually incorrect. In general, using the RDP classifier with the CO1 Eukaryote v1 training set and the recommended minimum bootstrap support cutoff at the genus rank significantly reduces the FPR.

**Table 4:** Taxonomic assignment outcomes at the genus rank from primer-anchored 200 bp sequences using the top BLAST hit method compared with the RDP classifier with the Eukaryote CO1 v1 training set.

| **Method** | **N\*** | **No results returned \*\*** | **TP** | **FN** | **TN** | **FP** | **Accuracy** | **TPR** | **FPR** |
| --- | --- | --- | --- | --- | --- | --- | --- | --- | --- |
| Top BLAST hit approach | 17,960,965 | 1,642 | 17,559,411 | 3,350 | 384 | 397,820 | 98% | ~100% | ~100% |
| RDP Classifier Eukaryote CO1v1 | 17,962,607 | N/A | 16,887,619 | 727,269 | 230,262 | 117,457 | 95% | 96% | 34% |

TP = true positive
FN = false negative
TN = true negative
FP = false positive
TPR = true positive rate
FPR = false positive rate
~ Indicates that the value was rounded up and is nearly 100%
\*N = Total number of primer-anchored 200 bp CO1 sequences used as queries during leave-one-out testing
\*\*BLAST results were not returned because the expect value was greater than 10

We also compared the time needed to make high-throughput sequence-based taxonomic assignments using the top BLAST hit method and the RDP classifier (Figure 4). Using a single processor, making assignments using the RDP classifier with the CO1 Eukaryote v1 training set was on average ∽19 times faster than using the top BLAST hit method. We did not consider the extra time needed to process tabular BLAST output into a usable format by adding taxonomic lineages and calculating query coverage.

**Figure 4:**
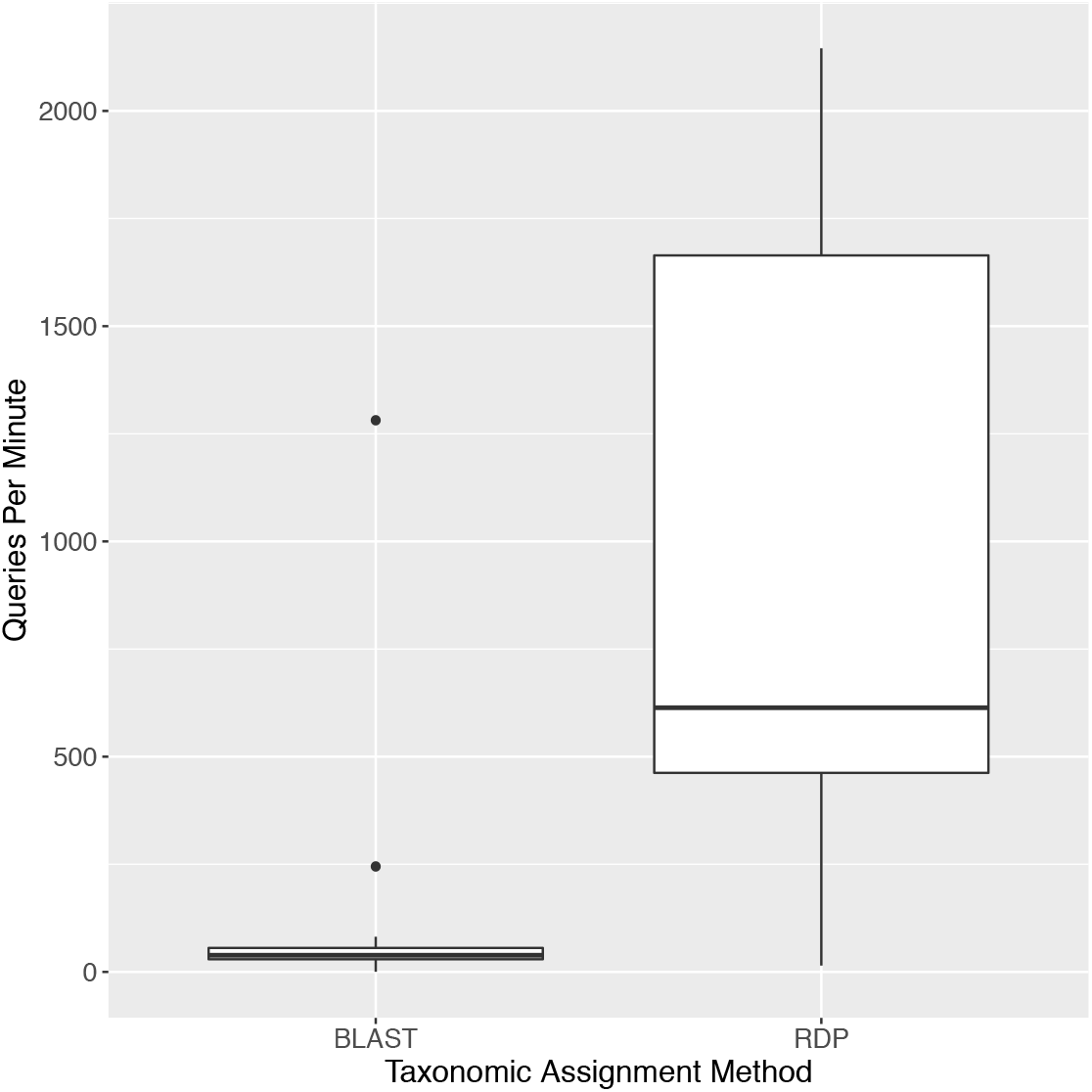
The RDP classifier taxonomically assigns more queries per minute than the top BLAST hit method. The number of primer-anchored 200 bp query sequences taxonomically assigned per minute is compred using the top BLAST hit method against a locally installed copy of the nucleotide database and the RDP classifier 2.12 with the CO1 Eukaryote v1 training set.

Compared with a 2013 training set, we found ∽3 times more class Insecta reference sequences 500 bp+ identified to the species rank (561,841 versus 190,333) and one additional order ‘Zoraptera’ from GenBank (Table S10). The group of top five Insecta orders with the greatest number of reference sequences has not changed from 2013 to 2016, though each group contains many more reference sequences today than 3 years ago (Table S11). The bottom five Insecta orders with the least number of reference sequences has changed slightly from 2013 and in the current training set includes Grylloblattodea (n=1), Zoraptera (n=2), undef_Insecta (n=9), Mantophasmatodea (n=30), and Dermaptera (n=37) (Table S12). As expected, as the number of reference sequences in the database grows, the proportion of genus rank incorrect assignments decreases (Table S13). Representation of various Insecta orders is shown in detail for CO1 sequences in the large class Insecta (Table S8). To further reduce misclassification rates in class Insecta, reference sequences for the Grylloblattodea, Zoraptera, Lepidotrichidae, Lepismatidae, Nicoletiidae, Mantophasmatodea, and Dermaptera need to be added to public databases.

## Discussion

CO1 metabarcoding has been extensively compared with morphology-based biomonitoring methods across a range of applications and has been repeatedly shown to detect more and/or a complementary suite of taxa compared with traditional methods ^13^. The continued interest and growing popularity of DNA metabarcoding in a diverse array of fields such as forestry, agriculture, fisheries, biosecurity, and conservation is driven by the scalability of this method when coupled with high throughput DNA sequencing ^14^. Efficient detection of taxa important for biomonitoring in particular relies on standardized, representative, and reproducible field sampling methods such as those developed by the Canadian Aquatic Biomonitoring Network (CABIN) or the Australian River Assessment Scheme (AUSRIVAS) ^15, 16^. Improvement of lab methods, such as primer development for PCR, and the use of multiple markers to increase detection coverage, and the development of PCR-free methods is also an active area of research ^17–20^. One major bottleneck in these efforts has been at the bioinformatics step and the ability to provide *high throughput* and accurate taxonomic assignments for CO1 metabarcodes using a purpose-built classifier, which is where our method fits in to this scheme.

Developments in the field of CO1 taxonomic assignment are mostly geared to the assignment of single queries though some methods can assign batches of sequences at once. Methods range from tools that use HMM alignment followed by a linear search ^11^, Neighbor-joining analysis ^1^, BLAST ^21^, minimum distance and fuzzy set theory ^22^, the coalescent ^23^, segregating sites ^24^, neural networks ^25^, and support vector machines ^26^. Other than the first three methods, none of the alternative methods have caught on for CO1 taxonomic assignment most likely because the average user is not aware they exist and there is no portal to allow for easy implementation. Our method leverages the well-known RDP Classifier which uses a naïve Bayesian method to taxonomically assign prokaryote 16S rDNA as well as fungal ITS and LSU rDNA and adapts it for use to classify animal CO1 mtDNA. Aside from capabilities demonstrated in this study, we believe the long history of this method, existing portal, open-source availability, and clear documentation will help widespread applicability of this method for fast and accurate high throughput taxonomic assignments.

To date, the most commonly used method for high throughput CO1 taxonomic assignment is the top BLAST hit method. Unfortunately, the BLAST metrics commonly used for delimiting good taxonomic assignments such as % identity, query coverage, bit score, e-value, or combinations thereof simply provides different measures of similarity to a top hit and a measure of random background noise in the database ^27^. The RDP classifier, on the other hand, was developed specifically to make taxonomic assignments from marker gene sequences and provide a measure of confidence to assess how likely the assignment is to be correct ^3^. With the top BLAST hit method, if there is no top BLAST hit that meets the user's criteria for a good assignment, then no assignment can be made. With the RDP classifier, if there are no congenerics in the database or if the genus rank assignment has a low confidence score, it may still be possible to make an assignment to a more inclusive rank if there are, for example, confamilial sequences in the database. In this study, we provide a matrix of minimum bootstrap support values to accommodate a range of CO1 sequence lengths and taxonomic assignment ranks that should provide 99% correct assignments assuming the CO1 query sequences are present in the training set, an implicit assumption for nearly every taxonomic assignment method. We show here that the RDP classifier is able to significantly reduce false positive rates compared with BLAST by using bootstrap support values at each rank as a filter for high confidence assignments.

The impact of different kinds of taxonomic assignment errors has been discussed in the literature ^28^. In this study, a false positive was defined as a sequence taxonomically assigned with high confidence even though it is wrong. Type I error also encompasses this outcome and generally refers to the incorrect rejection of a true null hypothesis. This is especially significant when the cost of making a misidentification is high, such as when a false positive assignment leads investigators to an over-estimation of the presence or distribution of a rare threatened or endangered species ^29^ or the assignment may create false alarm for an invasive or harmful species. In such cases, the RDP classifier is a more reliable tool to use than BLAST.

In this study a false negative was defined as a sequence correctly classified but with a confidence score below the threshold cutoff. Type II error also encompasses this outcome but generally refers to incorrectly retaining a false null hypothesis, i.e. when a sequence cannot be classified because congeneric sequences are missing from the database. This latter scenario is of particular relevance when missing the detection of a taxon of interest, such as in a quarantine situation could result in the introduction of parasites, pathogens, or invasive species ^28, 30^. In theory, in a quarantine situation where a limited suite of taxa is of interest, it should be easier to compile a representative database. The RDP classifier is more prone to FN's than BLAST, but as representative databases grow, high confidence assignments should improve by reducing the rate of false negatives due to missing congenerics in the database.

Previous work has shown that as few as 12% of described extant Insecta genera (8,679 / 72,618) are currently represented by full length (500 bp+) CO1 sequences identified to the species rank in the GenBank nucleotide database ^12^. Querying the database three years later, we found out that 22% of extant Insecta genera (16,285 / 72,618) are now represented by full length sequences identified to the species rank in the GenBank nucleotide database. At this rate of growth it could take 27 more years for all extant Insecta genera to be represented by a full length CO1 sequence in GenBank. There are about twice as many sequences available in BOLD compared with what is currently publically available in GenBank. This is only the tip of the CO1 barcode iceberg as the number of insect species is exponentially higher than genera and CO1 sequence representation in databases is expected to be even less than at the genus rank. This data gap could have significant implications to leverage the full potential of CO1 metabarcoding in current studies. In this study, we show a reduction of incorrect taxonomic assignments for major Insecta orders as databases have grown over the course of 3 years. We suggest that an immediate way to improve the number of high confidence assignments for current studies is to sequence CO1 barcodes for common representatives of local fauna to supplement existing databases.

## Methods

Three sets of CO1 reference sequences were assembled: 1) Arthropoda, 2) Chordata, and 3) outgroup taxa as described below using Perl with BioPerl modules and the Ebot script ^31, 32^. The following search terms were used to query the NCBI taxonomy database: 1) “Arthropoda”[ORGN] AND “species”[RANK] [Aug. 10, 2016], 2) “Chordata”[ORGN] AND “species”[RANK] [Aug. 24, 2016], and 3) “cellular organisms”[ORGN] AND “species”[RANK] NOT (“Arthropoda”[ORGN] OR “Chordata”[ORGN]) [Oct. 24, 2016]. A formatted taxon list was created using only taxa with complete binomial species names excluding the names containing sp., nr., aff., and cf. The NCBI nucleotide database was queried using the Entrez search term “cox1[gene] OR coxI[gene] OR CO1[gene] OR COI[gene] AND” the formatted taxon lists from above. For the outgroup taxa, the additional term “BARCODE”[keyword] was used. Sequences were retained if they were at least 500 bp and multiple sequences per species were retained when available. The associated taxonomic lineage was retrieved for each sequence. Human contaminant sequences were identified using BLAST and a custom database comprised of only human CO1 sequences. The taxonomic reports of hits with high query length coverage and high percent identity to known human sequences were individually explored, removed where necessary, and reported to NCBI. The Arthropoda, Chordata, and outgroup taxa were combined to create the CO1 Eukaryote v1 set trained to the genus rank and used with the RDP classifier v 2.12 for leave-one-out testing, cross-validation testing, and classifier training. A CO1 Eukaryote v2 training set was also created using the same sequences from above but was trained to the species rank.

Since metabarcoding samples often contain partially degraded eDNAs, shorter fragments are often targeted to increase PCR and sequencing success. As a result, leave-one-out testing was completed for full length (500bp+) CO1 sequences as well as for 400bp, 200bp, 100bp, and 50bp fragments. During leave-one-out testing, a sequence is removed from the dataset before it is classified. An assignment is scored as correct if the assignment matches the known taxonomy for the sequence. This assignment is made using a full set of 8 bp ‘words’ subsampled from the query sequence. Bootstrap support is assessed by subsampling a portion of the 8 bp ‘words’ from the query sequence, making an assignment, and counting the proportion of times the original taxonomic assignment is recovered. This is repeated 100 times. The sequence is returned to the training set and the next sequence is removed, classified, and so on. The purpose of this type of testing is to assess classifier performance (see below).

CO1 primers from the literature, especially those targeting invertebrates or developed especially for metabarcoding eDNA were compiled. Primers tested in this study and their references are shown in Table S14. These primers were aligned against the *Drosophila yakuba* CO1 region obtained from GenBank accession X03240 using Mesquite v 3.10 ^33^. CO1 secondary structure features from *Bos taurus* were obtained from UniProt accession P00396. We used CUTADAPT v1.10 to retrieve primer-trimmed sequences using default settings (allowing up to a 10% mismatch in the primer sequence) from our CO1 training set in the same way that real raw sequence data would be processed with the default settings ^34^. These sequences were trimmed to 200 bp fragments to simulate the average length of an Illumina read after primer trimming and we tested assignment accuracy and coverage using leave-one-out and cross-validation testing. For each primer, the RDP classifier was directly compared with the top BLAST hit method for taxonomic assignment. Assignments were compared at the genus rank for each method. ‘Good’ assignments for the RDP classifier was defined according to Table 2 for 200 bp fragments at the genus rank, requiring a bootstrap proportion of 0.60 or greater. ‘Good’ assignments for the top BLAST hit method was defined by having a top BLAST hit with percent identity >= 95% and a top BLAST hit alignment that spans >=85% of the original query sequence length (query coverage). We measured the proportion occurrence and rate of different types of taxonomic assignment outcomes as defined in Figure S5.

We also compared how class Insecta sequence database composition and incorrect taxonomic assignment distribution across insect orders have changed over the past three years. This was done by comparing the proportion of incorrect assignments from class Insecta in the current CO1 Eukaryote v1 training set [August 2016] with the Insecta Genbank-Genus training set [March 2013] that both used the leave-one-out testing method provided by the RDP classifier tool ^12^.

## Data availability

The trained data to be used with the RDP classifier are available for CO1 Eukaryote v1 (trained to the genus rank) and CO1 Eukaryote v2 (trained to the species rank) as supporting online material. The taxonomy and fasta files used for training are available from the corresponding author upon request.

## Acknowledgements

We would like to acknowledge funding for T. Porter from the Government of Canada through the Genomics Research Development Initiative as well as office space and computational resources provided by the Hajibabaei lab at Centre for Biodiversity Genomics, University of Guelph.

## Author contributions

T. Porter conceived of the manuscript idea and conducted the analyses. T. Porter and M. Hajibabaei wrote the manuscript.

## Additional information

The authors declare no competing financial interests.

Requests for materials and correspondence can be addressed to T.M. Porter at.

